# Ventral Striatal Dopamine Increases following Hippocampal Sharp-Wave Ripples

**DOI:** 10.1101/2025.07.24.666687

**Authors:** Miriam A. Janssen, Hung-tu Chen, Nicolas X Tritsch, Matthijs A. A. van der Meer

## Abstract

The reactivation of task-related hippocampal activity during sharp-wave ripples (SWRs), often expressed as sequential “replay,” is thought to contribute to behavior through both online and offline processes. During sleep and awake rest, replay supports memory consolidation, broadly defined as the updating and stabilization of knowledge structures for later use. During active task engagement, replay may also support decision-making through episodic memory retrieval and the generation of prospective scenarios^1–4^ (but see^5^). While disrupting SWR events impairs memory-guided behavior^6–9^ and augmenting them enhances such behavior^10,11^, the neurophysiological mechanisms that enable replay to contribute to behavior remain unclear. Theories of replay-based learning posit the need for replay events to be followed by an evaluative signal. Dopamine (DA) is consistently associated with such teaching signals, particularly in the ventral striatum, where it reflects reward- and other prediction-error signals^12^. Although hippocampal activity can influence mesolimbic dopamine signaling^13–15^, whether SWRs are coupled to ventral striatal DA release is unknown. To address this, we simultaneously recorded dorsal CA1 local field potentials to detect SWRs and measured ventral striatal DA concentration using fiber photometry of the GRAB_DA2m_ sensor^16^ in freely moving mice. We found a significant increase in DA following SWRs. This coupling was more prominent during offline rest than during periods of immobility during behavior, and was stronger following longer and higher-power SWRs. These results reveal coupling between hippocampal SWRs and striatal dopamine, providing a potential candidate mechanism through which internally generated hippocampal activity could engage evaluative signals during offline memory processing.

## Results

### Validating Dopamine Photometry with Amphetamine and Reward

We detected hippocampal sharp wave-ripples (SWRs, recorded from dorsal CA1 with custom microdrives) and measured dopamine (DA, using simultaneous fiber photometry) in the ventral striatum of wild-type C57BL6/J mice (n = 8; 5 male, 3 female) during rest periods before and after a reward-approach task designed to reveal putative reward prediction errors. After histological verification of electrode and fiber placement (Extended Data Figure 1), we sought to validate the fluorescent dopamine sensor GRAB_DA2m_^16^ and to calibrate DA changes relative to *bona fide* reward prediction errors evoked by the unexpected omission or delivery of rewards. To confirm that the measured fluorescence signal reflects DA, we first systemically administered amphetamine (AMPH), a dopamine reuptake inhibitor (Figure 1a). As expected, AMPH evoked a clear and persistent increase in ventral striatal DA compared to saline controls (baselined post-injection DA for AMPH compared to control sessions, paired, one-tailed Wilcoxon signed rank test: p = 0.0039; Figure 1c). We ran this same validation step on three mice expressing a green fluorescent protein insensitive to DA (“GFP-only” mice, n = 3; all male) to control for non-specific changes in fluorescence linked to changes in breathing, movement or hemodynamics. As expected, GFP-only baselined peak values in the post-injection incubation period were not significantly different between AMPH and control sessions (paired, one-tailed Wilcoxon Signed Rank test: p = 1; Figure 1c, inset).

**Figure 1.**
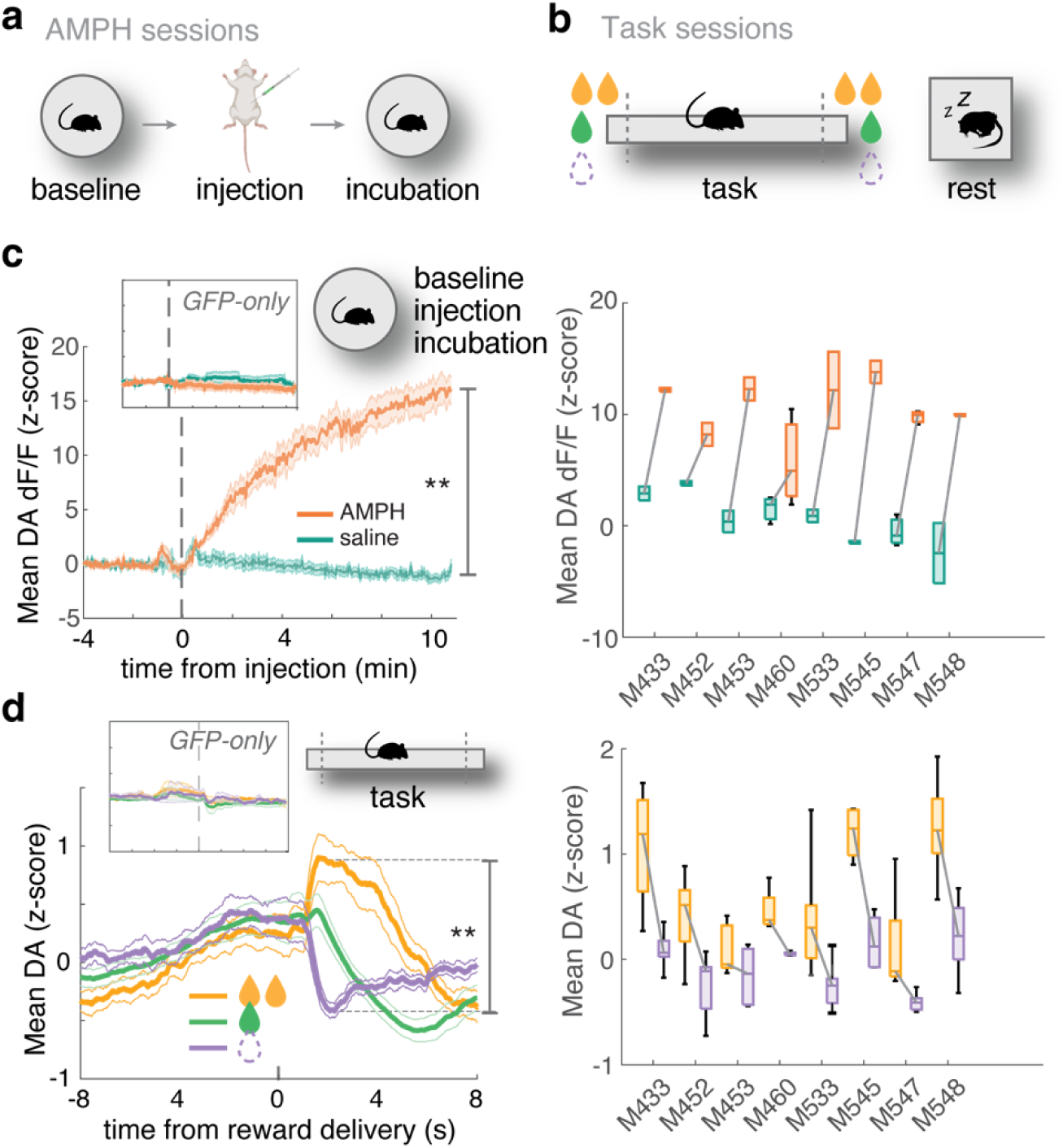
Ventral striatal DA tracks reward outcomes. **a,** Schematic of the experimental design for amphetamine injection sessions, showing a baseline rest period, intraperitoneal injection of amphetamine or saline, and an incubation rest period. **b**, Schematic of the experimental design for task sessions, consisting of a probabilistic reward approach task on a linear track and rest periods. The *rest* epochs occur before and after the linear track task to test if DA is coupled with SWRs during offline rest. The *task* epoch tests whether [DA] scales bidirectionally with reward: a positive reward prediction error occurs when animals receive 2x volume of evaporated milk reward at either end of the track (10 percent of trials; orange), no prediction error when animals receive expected 1x volume of reward (80 percent of trials; green), and a negative reward prediction error when animals receive no reward (10 percent of trials; purple). **c**, (left) average DA (z-scored dF/F) increased significantly following amphetamine (AMPH, 0.625 – 2.5 mg/kg, i.p.) compared to saline injection (paired Wilcoxon signed-rank test, p = 0.0039). Inset: average DA following AMPH (2.5 mg/kg) or saline in control mice expressing GFP only (paired, Wilcoxon signed-rank test, p = 1). Shaded region indicates ± SEM (*n* = 8 mice for GRAB_DA2m_, *n* = 3 for GFP-only). (right) Individual subject mean DA for AMPH (orange) vs saline (teal) injections; individual mice all had significantly larger DA for AMPH compared to saline (permutation test, all p < 0.05). **d**, (left) Putative reward prediction error signal in the average DA fiber trace (z-scored) aligned to reward zone entry, separated by reward magnitude: unexpected high reward (orange), expected medium sized reward (green), or unexpected omission of reward (purple). Paired, one-tailed Wilcoxon signed-rank test between omission and high-volume trials, p = 0.0039. Inset: absence of RPE signal in GFP-only mice (paired, one-tailed Wilcoxon signed-rank test between omission and high-volume trials, p = 0.125). Shaded region indicates ± SEM (*n* = 8 mice for GRAB_DA2m_, *n* = 3 for GFP-only). (right) mean DA within 4 seconds of a high-volume reward trial or omission trial photobeam break. All mice had numerically larger mean values for high-volume trials compared with omission trials, and 7 out of the 8 mice had significantly larger mean DA values for high-volume trials compared to omission trials (one-tailed permutation test all p < 0.05 except M453, p = 0.052). *** indicates p < 0.001, ** indicates p < 0.01, * indicates p < 0.05.

Next, to obtain putative reward prediction error (RPE) signals during behavior, we trained mice to perform a probabilistic reward delivery task, where rewards of different sizes were delivered at both ends of a linear track: medium-sized rewards (evaporated milk) on 80% of trials, large rewards (2x the medium reward) on 10% of trials, and no reward on the remaining 10% of trials (Figure 1b). Consistent with previous work ^17,18^, we found a clear suppression of the DA signal following reward omission relative to the expected (medium) amount, and a clear increase in DA following larger-than-expected reward (Figure 1d). Specifically, the high-volume reward trials had a mean DA value of 0.63 ± 0.17 z (SEM), the medium volume reward trials 0.15 ± 0.10 z, and the omission trials -0.067 ± 0.08 (0-4 s following reward zone entry, paired one-tailed Wilcoxon signed-rank test between omission and high-volume trials, p = 0.0039). GFP-only mice did not show any significant difference between mean DA values (paired, one-tailed Wilcoxon signed-rank test between omission and high-volume trials, p = 0.125, Figure 1d inset). Thus, together these observations demonstrate that we are measuring a putative DA RPE signal from the ventral striatum.

### Offline Ventral Striatal Dopamine Increases Following Hippocampal Sharp-Wave Ripples

Having established we could measure ventral striatal dopamine reflecting putative reward prediction errors, we next examined the relationship between hippocampal SWRs (total n = 24,253 SWRs detected across 92 rest sessions using our previously published frequency-spectrum similarity score, Carey et al. 2019 and *Methods*^19^; mean 267 ± 155 SWRs per session; see Extended Data Figure 2 for details; note that we use SWRs as a proxy for putative replay events and did not record spiking activity required for decoding these events) and ventral striatum DA during rest periods (Figure 2a). Apparent increases in DA following SWRs could be seen by the naked eye (see raw traces in Figure 2d). To quantify this effect, we generated SWR-triggered DA (SWR-DA) peri-event time histograms (PETHs) for each daily recording session. Averaging across sessions, we observed a clear increase in DA following SWRs (Figure 2b; median magnitude: 0.064 z ± 0.021 (SEM); peak time: 0.38 s ± 0.16 after detected SWRs). A session-based linear mixed effects regression model (LMM) with random intercepts for mice and baselined peak DA values as the dependent variable revealed a significantly positive intercept, indicating that the SWR-triggered DA peak was increased relative to the pre-SWR baseline peak (β = 0.079, t(181) = 3.37, p = 0.00091, CI: 0.033, 0.12; Figure 2b). We chose peak height because of our a priori hypothesis of a phasic dopamine transient, analogous to a reward prediction error, rather than slower changes; nevertheless, using baselined mean DA instead of peak DA similarly revealed a (borderline) significant post-SWR increase (β = 0.046, t(181) = 1.98, p = 0.050, CI: 5.39e-05, 0.092). Importantly, session-based LMMs on the GFP-only mice revealed no evidence for a SWR-triggered increase in peak DA (β = -0.00049, t(47) = -0.23, p = 0.82, CI: -0.0049, 0.0039) or mean DA (β = -0.0022, t(47) = -1.44, p = 0.16, CI: -0.0053, 0.00089). Additionally, we ran the same analysis with a more lenient SWR detection method (peak ripple-band power > 3 z, n = 67,865 SWRs). Again, SWR triggered peak DA was greater after a SWR compared to before (β = 0.067, t(181) = 4.28, p = 2.98e-05, CI: 0.036, 0.098) and SWR triggered mean DA response was similarly increased (β = 0.039, t(181) = 2.38, p = 0.018, CI: 0.0067, 0.072). To contextualize this result, we compared the size of the internally generated SWR-DA transients with the DA transients generated following the receipt of high-volume rewards (putative positive RPEs) during task performance. The size of the internally generated SWR-DA transient was 23.8% of the size of a putative positive RPE (the difference between unexpected high volume reward trials and expected medium volume reward trials).

**Figure 2.**
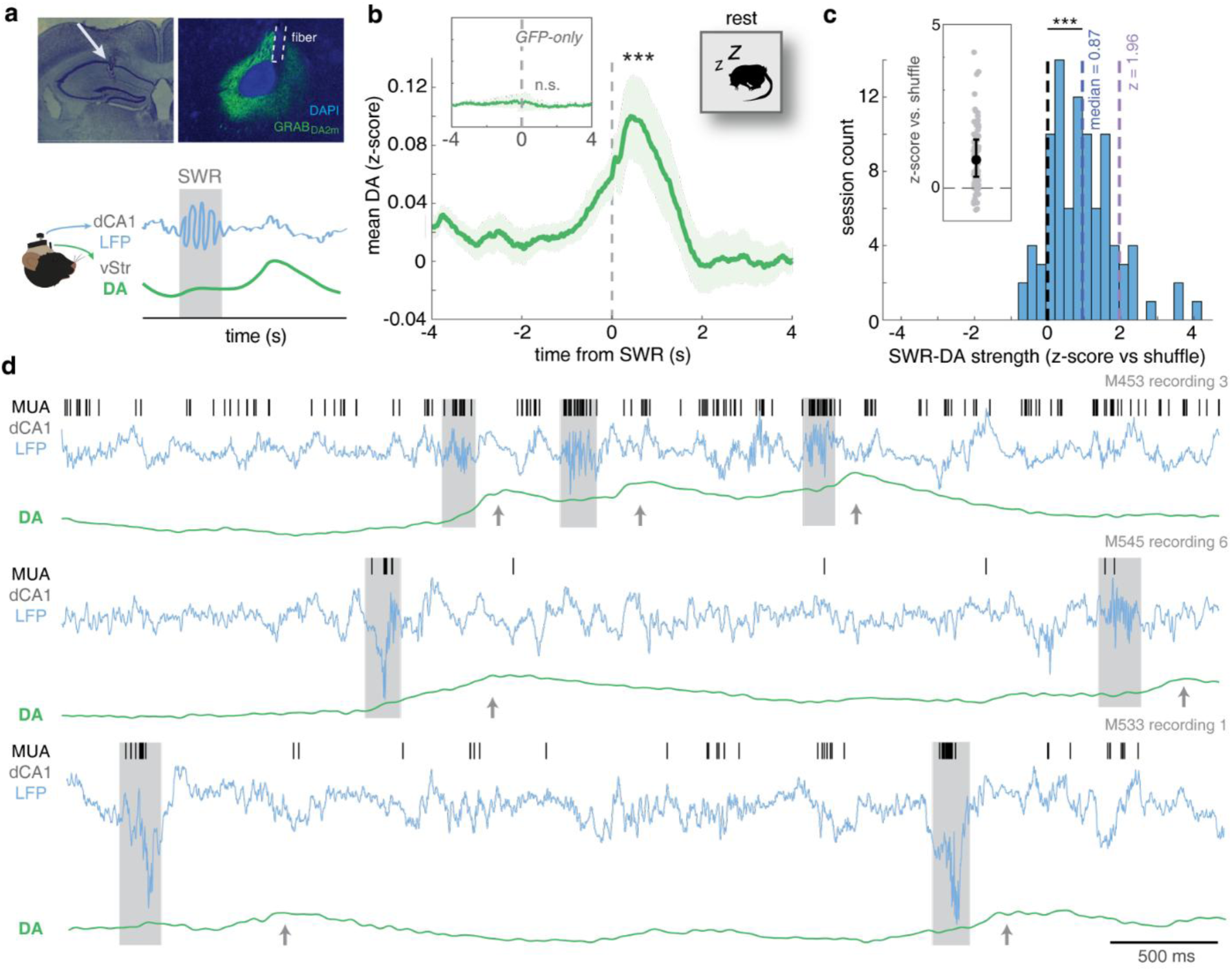
Ventral striatal dopamine increases following hippocampal sharp wave-ripples. **a**, (top) Example CA1 coronal sections indicating microelectrode placement (arrow) and simultaneous ventral striatal GRAB_DA2m_ expression for fiber photometry (DAPI, blue; GFP, green; see Extended Data Figure 1 for full histology) and fiber placement (dashed lines). (bottom) Schematic of hypothesized results: a transient increase in the ventral striatal fiber photometry dopamine signal (DA, green line) following dorsal CA1 sharp wave-ripples (SWR, blue line and shaded region). **b**, Ventral striatal DA increased significantly following SWRs (dashed vertical line at t = 0; peak DA 1 second after SWRs compared to before, LMM intercept, p < 0.001). Inset: as in b, but for GFP-only mice (n.s.). Shaded region indicates ± SEM (*n* = 8 mice for GRAB-DA, *n* = 3 mice for GFP-only). **c**, Distribution of SWR-DA coupling strength across recording sessions (n = 92 rest sessions; one pre-task and one post-task session per recording day), as computed by z-scoring relative to a resampled distribution where the DA signal was circularly shifted relative to SWR times. SWR-DA coupling across sessions was significantly greater than zero (median = 0.87 z, Wilcoxon signed-rank test. z-value = 7.49, p = 3.55 x 10^-14^). Inset: Same data replotted as filled circles for each session and errorbar indicating the 95% confidence interval. **d,** Example multiunit activity (MUA, black tick marks), local field potential traces (LFP, blue line) and GRAB_DA2m_ fiber photometry traces (detrended and z-scored DA signal, green line) for three sessions. Detected SWRs and corresponding MUA are shaded in grey. Following these SWRs, transients in the DA signal are visually apparent (grey arrows). *** indicates p < 0.001, ** indicates p < 0.01.

By inspecting individual sessions, we observed notable variability in the magnitude and timing of SWR-DA coupling (Extended Data Figure 3). To describe this variability, we computed single-session PETHs, which for each session was z-scored relative to a distribution of resampled PETHs in which the DA signal was circularly shifted by a random amount. Across sessions, median SWR-DA coupling was significantly greater than zero (Figure 2c; median = 0.87 z; Wilcoxon signed-rank test z-value = 7.49, p = 3.55 x 10^-14^; note 82 sessions showed positive z-scores). Out of 92 rest sessions (one pre-task rest and one post-task rest per recording day), 11 sessions had individually significantly positive associations (z > 1.96), while none had a negative association (z < -1.96; Figure 2c). This variability across sessions could not be explained by differences in running speed on the track (a proxy for motivation; likelihood ratio test between LMMs with and without running speed, LRStat_Δdf=1_ = 0.34, p = 0.58) or by the magnitude of putative reward prediction error coding during the task (a proxy for GRAB-DA expression and fiber location; likelihood ratio test, LRStat_Δdf=1_ = 0.19, p = 0.66). Thus, the distribution of SWR-DA coupling across sessions, while variable, is clearly shifted relative to 0, indicating that significance is not an artifact of a few sessions at the positive tail of a null distribution.

### SWR properties and behavioral state contribute to SWR-DA coupling

To investigate this variability with more granularity, we next sought to model not only the effects of training and task variables (testing day & pre- vs. post-task rest), but also the effects of SWR properties (power, length) and behavioral state (sleep vs. wake) using event-based, rather than session-based, linear mixed effects regression model comparisons. As before, we used baseline-corrected peak DA as the dependent variable, quantified for each individual SWR event as the peak DA signal during the 1s period following SWR onset minus the peak DA signal during the 1s period preceding SWR onset.

Consistent with the session-based model, the event-based LMM with random intercepts for mice and baselined peak DA values as the dependent variable revealed a significantly positive intercept, indicating that the average baseline-corrected SWR triggered DA response was increased relative to pre-SWR baseline (β = 0.085, t(24252) = 3.44, p = 0.00059, CI: 0.037, 0.13; Table 1). For completeness, we also ran a parallel set of models with baselined mean SWR-DA (Table 2), with comparable results (β = 0.051, t(24252) = 2.16, p = 0.031, CI: 0.0047, 0.097; Table 2). With these base models established, we next sought to determine the contributions of different task, behavioral, and SWR event properties. In the analyses that follow, we ask two complementary questions for each predictor: (1) whether adding that predictor to the base random-effects model improves model fit, and (2) whether removing that predictor from the full model reduces model fit. The first tests the predictor’s marginal association with SWR–DA, whereas the second tests its unique contribution after the effects of other predictors are accounted for.

**Table 1.** Baselined Peak DA Linear Mixed Effects Model results.

| Parameter | Int. | Coeff. | P | 95% CI | LRT stat \| P |
| --- | --- | --- | --- | --- | --- |
| **Base Model** | 0.085 |  | **0.00059** | 0.037, 0.13 |  |
| Base + Pre/Post (Pre) | 0.094 | -0.014 | 0.15 | -0.034, 0.0053 | 2.04 \| 0.16 |
| Base + Early/Late (Early) | 0.092 | -0.015 | 0.15 | -0.034, 0.0054 | 2.04 \| 0.16 |
| Base + Sleep (Wake) | 0.073 | 0.024 | **0.015** | 0.0047, 0.044 | **5.86 \| 0.020** |
| Base + SWR duration | 0.00096 | 1.22 | **1.97e-11** | 0.86, 1.58 | **44.93 \| 0.001** |
| Base + SWR power | -0.069 | 0.056 | **1.40e-24** | 0.045, 0.067 | **104.73 \| 0.001** |
| Base + SWR ID | 0.073 | 0.024 | 0.15 | -0.0085, 0.056 | 2.09 \| 0.015 |
| **Base Model for Track Mice** | 0.056 |  | **0.014** | 0.011, 0.10 |  |
| **Base + Rest/Track (Track)** | -0.059 | 0.12 | **1.94e-07** | 0.077, 0.17 | **27.08 \| 0.001** |
| **Base model for Track Sessions** | -0.029 |  | 0.30 | -0.083, 0.026 |  |
| **Base model for Rest Sessions** | 0.065 |  | **0.003** | 0.022, 0.11 |  |
| Base + Pre/Post (Pre) | 0.091 | -0.043 | **0.00049** | -0.068, -0.019 | **12.14 \| 0.001** |
| **Full Model** | -0.10 |  | **0.00074** | -0.16, -0.044 |  |
| Full model – Pre/Post | -0.10 |  | **0.0011** | -0.16, -0.041 | **16.49 \| 0.001** |
| Full model – Early/Late | -0.11 |  | **0.00043** | -0.17, -0.048 | 0.54 \| 0.49 |
| Full model – Sleep | -0.10 |  | **0.00085** | -0.16, -0.042 | 0.96 \| 0.35 |
| Full model – SWR duration | -0.073 |  | **0.017** | -0.13, -0.013 | **14.59 \| 0.001** |
| Full model – SWR power | -0.010 |  | 0.72 | -0.066, 0.045 | **72.08 \| 0.001** |
| Full model – SWR ID | -0.09 |  | **0.0036** | -0.15, -0.029 | **11.29 \| 0.003** |
| **Full Model for Track Mice** | -0.15 |  | **0.00035** | -0.23, -0.067 |  |
| Full model – Pre/Post | -0.060 |  | 0.10 | -0.13, 0.012 | **8.66 \| 0.004** |
| Full model – Rest/Track | 0.00074 |  | 0.98 | -0.063, 0.065 | **27.12 \| 0.001** |

Notes: Reference group is in parentheses. *p*-values < 0.05 are bolded.

**Table 2.** Baselined mean DA Linear Mixed Effects Model results.

| Parameter | Int. | Coeff. | P | 95% CI | LRT stat \| P |
| --- | --- | --- | --- | --- | --- |
| **Base Model** | 0.051 |  | **0.031** | 0.0047, 0.097 |  |
| Base + Pre/Post (Pre) | 0.060 | -0.015 | 0.063 | -0.031, 0.0008 | 3.45 \| 0.069 |
| Base + Early/Late (Early) | 0.055 | -0.009 | 0.29 | -0.025, 0.0074 | 1.14 \| 0.30 |
| Base + Sleep (Wake) | 0.039 | 0.024 | **0.0033** | 0.0081, 0.040 | **8.65 \| 0.003** |
| Base + SWR duration | -0.016 | 0.96 | **8.94e-11** | 0.67, 1.25 | **41.99 \| 0.001** |
| Base + SWR power | -0.072 | 0.045 | **1.98e-23** | 0.036, 0.054 | **99.42 \| 0.001** |
| Base + SWR ID | 0.046 | 0.010 | 0.44 | -0.016, 0.037 | 0.59 \| 0.43 |
| **Base Model for Track Mice** | 0.022 |  | 0.33 | -0.022, 0.066 |  |
| Base + Rest/Track (Track) | -0.078 | 0.11 | **5.87e-09** | 0.071 0.14 | **33.86 \| 0.001** |
| **Base Model for Track Sessions** | -0.094 |  | 0.16 | -0.24, 0.049 |  |
| **Base Model for Rest Sessions** | 0.030 |  | 0.23 | -0.012, 0.071 |  |
| Base + Pre/Post (Pre) | 0.052 | -0.037 | **0.00011** | -0.056, -0.018 | **15.01 \| 0.001** |
| **Full Model** | -0.099 |  | **0.00028** | -0.15, -0.046 |  |
| Full model – Pre/Post | -0.096 |  | **0.00038** | -0.15, -0.043 | **14.86 \| 0.001** |
| Full model – Early/Late | -0.10 |  | **0.00019** | -0.15, -0.048 | **0.16 \| 0.74** |
| Full model – Sleep | -0.095 |  | **0.00036** | -0.15, -0.043 | 2.70 \| 0.097 |
| Full model – SWR duration | -0.074 |  | **0.0058** | -0.13, -0.021 | **13.60 \| 0.001** |
| Full model – SWR power | -0.024 |  | 0.36 | -0.076, 0.028 | **68.72 \| 0.001** |
| Full model – SWR ID | -0.089 |  | 0.00097 | -0.14, -0.036 | **7.51 \| 0.009** |
| **Full Model for Track Mice** | -0.16 |  | **6.31e-06** | -0.23, -0.090 |  |
| Full model – Pre/Post | -0.069 |  | 0.028 | -0.13, -0.0077 | **14.59 \| 0.001** |
| Full model – Rest/Track | -0.028 |  | 0.33 | -0.085, 0.028 | **32.56 \| 0.001** |

*Notes: Reference group is in parentheses. p-values < 0.05 are bolded.*

First, to address the contribution of elapsed time on SWR-DA coupling, we found the base model was not improved by distinguishing pre- vs. post-track rest (likelihood ratio test, LRStat_Δdf=1_ = 2.04, p = 0.16; Figure 3a). However, comparing a full LMM (see STAR methods) containing all variables and the same model without pre- and post-track rest labels did reveal a significant difference (likelihood ratio test, LRStat_Δdf=1_ = 16.49, p = 0.001). This was also the case with mean DA as the dependent variable (against base LMM: likelihood ratio test, LRStat_Δdf=1_ = 3.45, p = 0.069; against full LMM: likelihood ratio test, LRStat_Δdf=1_ = 14.86, p = 0.001), suggesting that a potential effect of pre- vs. post-track rest is only visible after the contributions of covariates are unmasked. Regression coefficients for both models indicated that pre-track rest SWR-DA coupling tended to be higher than during post-track rest (Figure 3a), as would be expected from gradual photobleaching of the GRAB-DA sensor. Contrary to this expectation, however, there was some indication that within a session, SWR-DA coupling increased with time when the SWR occurred (SWR_ID; n.s. relative to the base model, but a significant predictor in the full model: likelihood ratio test, LRStat_Δdf=1_ = 11.29, p = 0.003). Thus, multiple different processes likely contribute to the effects of time on SWR-DA coupling, although covariance with other variables prevent unambiguous conclusions.

**Figure 3.**
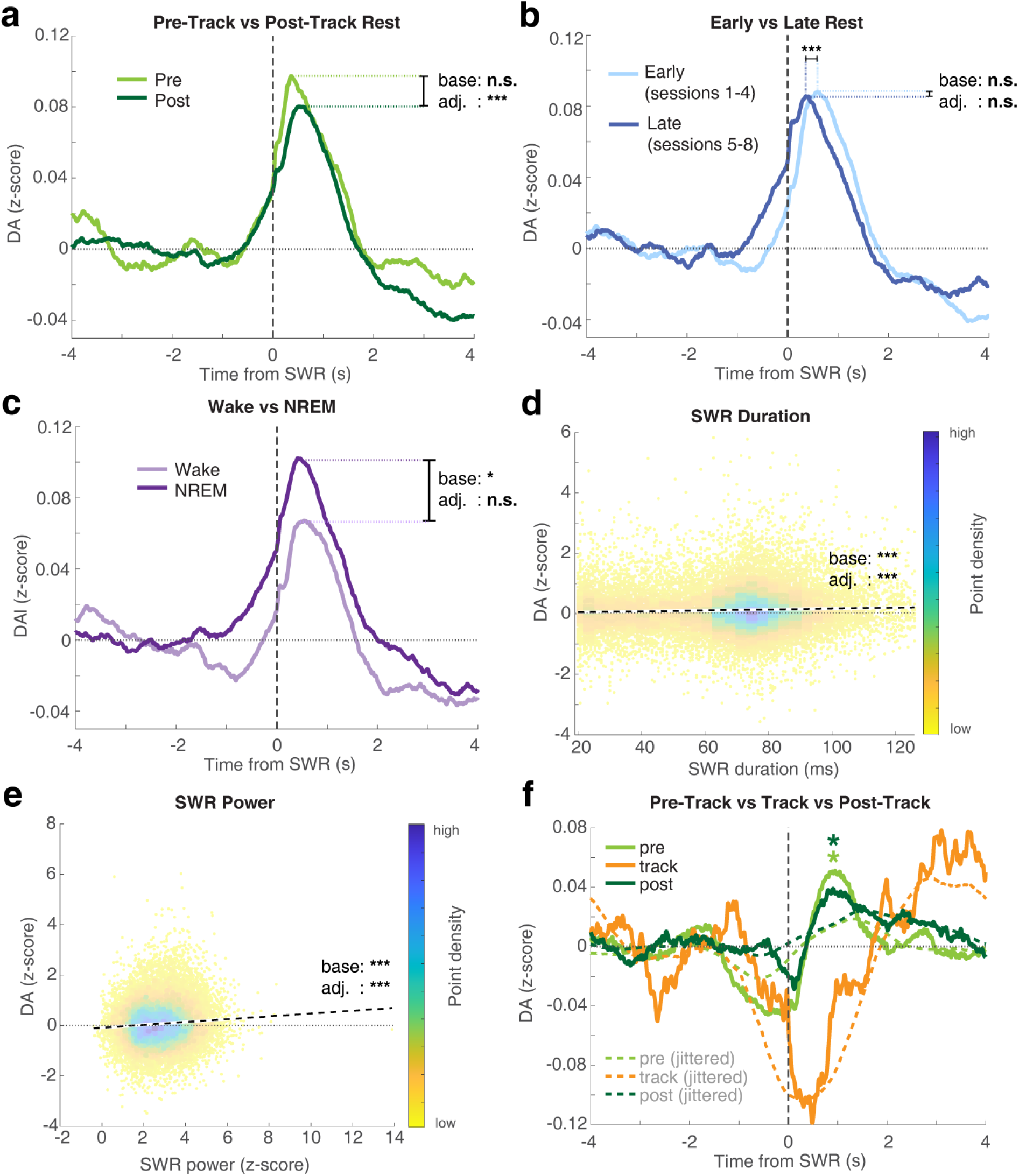
Event-by-event analysis of features contributing to SWR-DA coupling. **a,** SWR–DA coupling showed no clear overall pre/post-task difference (pre: n = 8,612 SWRs; post: n = 15,641 SWRs; base LMM model comparison n.s.), although a pre/post effect emerged after other predictors were included in the model (see main text); significance is reported for comparison of the base model with the predictor of interest added (‘base’) and for the comparison of the full model with that predictor removed (adjusted, ‘adj.’). **b**, SWR–DA coupling did not vary in strength between early (sessions 1–4; 46 sessions, n = 14,091 SWRs) and late sessions (sessions 5–8; 46 sessions, n = 10,162 SWRs). However, the SWR-DA peak shifted significantly earlier in late compared to early sessions (see main text). **c,** SWR–DA coupling during NREM sleep was significantly stronger compared to wakeful rest periods (wake: n = 12,207 SWRs; NREM: n = 12,046 SWRs), although sleep stage did not explain unique variance after adjustment for other covariates (see main text). **d,** SWR-DA peak value was significantly correlated with SWR duration (likelihood ratio test, p < 0.001 for both base and adjusted models, n = 24,253). Each point color indicates event density estimated from a two-dimensional histogram; dashed black line indicates line fit. **e,** SWR-DA peak value was significantly correlated with SWR power (likelihood ratio test, p < 0.001 for both base and adjusted models, n = 24,253). **f,** Replication of significant SWR-DA coupling during pre-track (light green line, n = 10,055 SWRs, intercept β = 0.079, p = 0.022) and post-track rest (dark green line, n = 14,496 SWRs, intercept β = 0.045, p = 0.029). SWR-DA coupling differed significantly between offline rest periods of immobility and periods of immobility on the track (orange line; likelihood ratio test, p < 0.001) and on-track rest did not show significant SWR-DA coupling (see main text). To distinguish slow task-related changes in the SWR-DA PETH from true fine-timescale coupling, we obtained jittered versions of each of the above PETHs (dashed lines) by circularly shifted the SWR-DA signal by a random time ± 1 second for each SWR to break the short-timescale coupling (averaged over 1,000 repeats). Note that this jittering reduces the SWR-DA peak for the pre- and post-task rest periods (dashed green lines), but not the slow changes on the task (dashed orange line). Significance bars indicate comparison between baselined peak values. *** indicates p < 0.001, ** indicates p < 0.01, * indicates p < 0.05.

Next, repeating this analysis with early recording sessions (days 1-4) vs. late sessions (days 5-8; note that mice had experience with the task prior to recording, so this is not a test of learning per se) recording did not significantly affect either the base model nor the full model (against base LMM: likelihood ratio test, LRStat_Δdf=1_ = 2.04, p = 0.16; against full model: likelihood ratio test, LRStat_Δdf=1_ = 0.54, p = 0.49; Figure 3b). This is in agreement with the mean DA LMM analysis (against base LMM: likelihood ratio test, LRStat_Δdf=1_ = 1.14, p = 0.30; against full LMM: likelihood ratio test, LRStat_Δdf=1_ = 0.16, p = 0.74). We did note a shift in the timing of the post-SWR DA peak, which moved earlier (closer to SWR time) for late sessions compared to early sessions (early peak time: 0.60 ± 0.31s following the SWR; late: 0.41 ± 0.29 (Figure 3b); LMM likelihood ratio tests all p < 0.01 compared to base model and full model; note these event-based peak time estimates differ slightly from the session-based estimate in Figure 2b).

If SWR-DA coupling is a potential mechanism for offline consolidation, one would expect larger SWR-DA transients during NREM sleep compared with periods of wakefulness. To test this, we ran event-based linear mixed effects model comparisons as above. The base model was improved by adding wake/NREM as a variable (likelihood ratio test, LRStat_Δdf=1_ = 5.86, p = 0.020), with the SWR-DA peak higher for NREM sleep (NREM beta with wake as the reference category: 0.024). Sleep stage predictiveness could, however, be explained by covariance with other variables (likelihood ratio test, LRStat_Δdf=1_ = 0.96, p = 0.35 when removed from the full model). This was in agreement with the mean (rather than peak) LMM (against base model: likelihood ratio test, LRStat_Δdf=1_ = 8.65 p = 0.003; against full model: likelihood ratio test, LRStat_Δdf=1_ = 2.70, p = 0.10). Session-by-session differences in the relative time spent in NREM sleep vs. rest did not explain significant variance either in the base model or when removed from the model, and were not significantly correlated with session-by-session measures of SWR-DA strength (z-scored relative to shuffle as in Figure 2b). Thus, the effect of wake vs. NREM sleep seems to be marginal rather than primary.

Finally, given that longer SWR duration length and higher SWR amplitude are associated with stronger PFC and CA1 reactivations and work showing prolonged SWRs enhanced behavioral performance^11,20^, we tested whether SWR properties duration and power are predictive of SWR-DA coupling magnitude. As before, we ran event-based linear mixed effects regression model comparisons with baselined peak SWR-DA values as the dependent variable. This model was improved by adding SWR duration and also SWR power as a variable in the model (likelihood ratio test, duration: LRStat_Δdf=1_ = 44.93, p = 0.001, power: LRStat_Δdf=1_ = 104.73, p < 0.001) (Figure 3d, e). Both SWR power and SWR duration could not be explained by their covariance with other variables (likelihood ratio test, duration: LRStat_Δdf=1_ = 14.59, p = 0.001, power: LRStat_Δdf=1_ = 72.08, p < 0.001). This was also in agreement with the mean DA LMM analysis results (against base model and full model, all p < 0.001). Overall, the robustness and magnitude of the model comparison differences and their t-statistics in the models mark out SWR power and duration as the strongest predictors of SWR-DA coupling among all variables we considered.

### SWR-DA coupling is stronger during offline rest compared to periods of on-task immobility

To determine if the SWR-DA relationship persists during active task engagement, we ran another cohort of wild-type C57BL6/J mice (n = 4; 3 female) on the same task. Instead of custom microdrives, we implanted 16-channel single-shank laminar silicon probes (NeuroNexus) that were light enough for the mice to move around the track while also tethered to the fiber. Each mouse was run for 8 recording days resulting in a total of 96 sessions, running 48.8 ± 5.1 trials on the track per session on average. To make a fair comparison between on-line and off-line SWRs, we only included SWRs during periods of immobility (314 ± 71 SWRs per session during pre-track rest, 57 ± 30 SWRs on the track, and 453 ± 95 SWRs in post-track rest). We first replicated our main result of SWR-DA coupling, using the event-based LMM analysis method with baselined peak DA as the dependent variable. As before, the SWR-DA intercept was significantly positive (post-SWR DA peak minus pre-SWR DA peak; intercept β = 0.056, t(26382) = 2.45, p = 0.014, CI: 0.011 0.10; Figure 3f), indicating a post-SWR increase in DA. As in our main data set, pre-task SWR-DA coupling was higher than post-task, which in this replication did reach statistical significance (likelihood ratio test vs. base model, LRStat_Δdf=1_ = 12.14 p < 0.001) which could not be explained by their covariance with other variables (likelihood ratio test vs. full model, LRStat_Δdf=1_ = 8.66, p = 0.004). This was also the case for the mean (rather than peak) LMM (against base LMM: likelihood ratio test, LRStat_Δdf=1_ = 15.01, p < 0.01; against full LMM: likelihood ratio test, LRStat_Δdf=1_ = 14.59, p < 0.001).

This replication of SWR-DA coupling during periods of immobility during offline rest is in contrast to what we found during periods of immobility on the task, where the baseline-corrected SWR triggered DA response was not significantly greater than zero, even trending negative (β = -0.047, t(1832) = - 1.75, p = 0.081, CI: -0.10, 0.0057; Figure 3f). This is in agreement with the baselined mean DA analysis (β = -0.094, t(1832) = -1.29, p = 0.20, CI: -0.24, 0.049). We noted that the on-track SWR-DA PETHs appeared to show a different temporal profile than the offline rest PETHs, including an apparent dip around SWR time t = 0, followed by a slow rise from 2-4 seconds. We hypothesized this profile is a consequence of the times that on-task SWRs tend to occur relative to behavioral states and task events known to produce DA changes. In particular, SWRs are likely to occur during inter-trial interval rest, when a reward-induced DA increase has recently occurred, and DA is likely to increase as animals start the next trial. Thus, to determine how much of the observed on-track SWR-DA PETH was due to such SWR-task correlations, we jittered each SWR by a random time between [-1 1] s, thus preserving associations with slow behavioral events but breaking any fine-timescale coordination between SWRs and DA. The jittered PETH for on-task SWR-DA preserved the major features of its temporal profile, including the prominent dip around t = 0, and the slow rise from 2-4 s (Figure 3f, dashed orange line), demonstrating that this shape is not a consequence of direct SWR-DA coordination but likely results from slower correlations with task events. In contrast, the post-SWR DA peak for the pre- and post-task rest periods (at 0.95 and 0.91 s respectively) was significantly reduced by the jittering procedure (light green and dark green dashed lines, Figure 3f; observed SWR-DA vs. resampled jittered distribution, p < 0.001), indicating that SWR-DA coupling during rest reflects genuine fine-timescale coordination.

Finally, to test explicitly whether SWR-DA coupling differs between offline rest and on-task pauses, we ran an LMM with rest session and track session labels. A variable indicating session-type (rest vs. track) significantly improved the base peak model (likelihood ratio test, LRStat_Δdf=1_ = 27.08, p < 0.001, and the predictiveness could not be explained by their covariance with other variables (likelihood ratio test LRStat_Δdf=1_ = 27.12, p < 0.001). This was also the case for the mean DA LMM (against base LMM: likelihood ratio test, LRStat_Δdf=1_ = 33.86, p < 0.001; against full LMM: likelihood ratio test, LRStat_Δdf=1_ = 32.56, p < 0.001). As a final direct test, we ran direct comparisons of SWR-DA coupling, and found that pre-track rest and track session baselined peak SWR-DA values were significantly different (Wilcoxon rank sum test, p = 1.97e-05), post-track rest and track session baselined peak SWR-DA values were significantly different (Wilcoxon rank sum test, p = 5.57e-06), and pre- and post-track rest session baselined peak SWR-DA values were not significantly different (Wilcoxon rank sum test, p = 0.31). Thus, SWR-DA coupling was significantly stronger during offline rest (including pre-task rest) compared to on-task immobility, and we did not find evidence for SWR-DA coupling on the task.

## Discussion

This study establishes for the first time a link between sharp wave-ripples (SWRs; putative replay events) in CA1 and dopamine transients in the ventral striatum, a downstream projection target of the hippocampus and a major node in brain networks that support reinforcement learning and decision-making. Prominent theories on how hippocampal replay influences subsequent behavior postulate a link between replay events and an evaluative signal that can drive learning and/or decision-making. For instance, Sutton et al. (1991) and Lin et al. (1992) used replay to generate fictive reward prediction errors which in turn trigger updating of learned values that ultimately drive behavior^21,22^. This offline learning is notable as it has the potential to solve the temporal credit assignment problem (see Mattar & Daw 2018; Sagiv et al. 2025 for influential applications to hippocampal replay^23,24^). A different proposal for the function of replay, that it may guide on-line decisions through retrieval of relevant memories and/or mental simulation of possible futures^4,25^ (but see van der Meer & Bendor 2025^5^), similarly requires a downstream evaluative step to translate memory into behavior. Our results lend new experimental support to these theories by demonstrating that hippocampal SWRs are followed by transient increases in ventral striatal dopamine, an established focal point for learning, motivation, and goal-directed behavior^12^.

This work is congruent with, and extends, prior studies showing that CA1 SWRs modulate the activity of ventral striatal neurons^26,27^ and reward-responsive VTA neurons^28^. However, this prior work left open whether these SWR-coupled VTA neurons were dopaminergic, and more generally, whether the reported changes in spiking activity resulted in measurable changes in ventral striatal DA – a particularly important issue given that ventral striatal DA levels may be uncoupled from DA neuron spiking in the VTA^29^. Addressing these limitations, the present study examined extracellular DA in the ventral striatum directly using fiber photometry, after establishing that the same signal encodes a reward prediction error during on-line behavior. The overall working hypothesis that an evaluative signal coupled to replay could drive learning in the brain is causally supported by the demonstration that medial forebrain bundle stimulation triggered on reactivation of a specific place cell results in place preference for that cell’s place field^10^. Our observation of SWR-DA coupling is a potential component of how such learning could occur endogenously.

Several features of the observed SWR-DA coupling are worth noting. First, while the main effect was robust, as demonstrated by the replication across two independent cohorts of mice and the convergence of multiple different statistical approaches (session-based circular permutation test; linear mixed models across events), we also observed substantial variability across sessions and events. Although some of that variability could be explained by the length and power of SWR events, and by conditional effects of wake/sleep state and time within recording session, striking differences across sessions remained, even within the same subject, and could therefore not obviously be attributed to fiber placement or virus expression. One potential factor underlying this variability is that we were blind to the actual “content” of SWR events, which likely included diverse experiences as well as events that are not readily decodable^2,3,30^; it is possible that if we had access to decoded SWR trajectories, it would turn out that only some, but not others, are associated with dopamine increases. Another contributor to the observed variability may be that although the task was not fully predictable, it was no longer novel given that mice had prior experience on the task before recording began. Novel experience can preferentially bias subsequent hippocampal reactivation toward the recent experience, whereas replay in familiar environments can include remote representations^31–33^. Future work capable of decoding SWR content during novel vs. familiar experience could test these ideas.

A second notable feature of the work is that we consistently observed SWR-DA coupling during offline rest – in a “sleep box” distinct from the actual behavioral task – but not during inter-trial periods of immobility on the task. A potential methodological concern in this comparison is that, as expected, we recorded fewer SWR events during on-task rest compared to offline rest; thus, a null result for on-task SWR-DA could have been the result of insufficient statistical power. Arguing against this possibility, however, is that direct statistical comparisons of on-task vs offline SWR-DA coupling were sufficiently powered to show a difference. Thus, a conservative interpretation of the results is that offline SWR-DA coupling is stronger than on-task SWR-DA. At first glance this result seems more consistent with the interpretation that SWR-DA serves offline consolidation rather than on-task performance, but it should be noted that the task was not only highly familiar, but also unlikely to engage the sort of memory- and inference-based processing likely to engage the hippocampus^3,4^; thus, further work should determine to what extent SWR-DA coupling depends on task demands. Within offline rest, pre-task SWR-DA coupling was as strong, or stronger (numerically in the first experiment, statistically in our replication) compared to post-task SWR-DA. While this difference was modest, we nonetheless consider it notable, particularly considering the alternative hypothesis that experience on the task would be expected to generate at least some motivationally relevant experiences worth replaying (as seen in e.g. post-task increase in hippocampal replay^33,34^), with larger SWR-DA coupling as a result. Moreover, Gomperts et al. found stronger coupling between SWRs and VTA neuron activity during wake, rather than sleep; the opposite of what we see here in SWR-DA coupling. Here, too, the introduction of highly novel experiences would be informative in determining the experience-dependence of SWR-DA, which in the current results only manifested as a temporal shift between early and late sessions.

The temporal relationship between SWRs and DA (DA peak following the SWR by ∼0.3-0.4s; the apparent increase in DA prior to the SWR could be the result of imperfect SWR detection) suggests that the SWR causing the DA transient is more likely than the reverse, but a reasonable null hypothesis is that there is no direct connection – rather, SWRs and DA transients may both be modulated by shared slower rhythms. Arguing against this interpretation is the observation that SWR-DA coupling is significantly decreased by temporal jitter of +/- 1 s, and the generally opposing brain states that are thought to be conducive to SWRs (rest, sleep) as compared to dopamine increases (activating). Moreover, there are multiple known anatomical and functional pathways through which hippocampal activity could result in ventral striatal DA increases, including direct projections to presynaptic DA terminals^13^. Additionally, hippocampal output via the subiculum may influence VTA activity through polysynaptic loops involving the nucleus accumbens and potentially ventral pallidum^35^. Indirect HC– PFC–VTA pathways are also anatomically possible, although inactivation of the mPFC did not attenuate hippocampal stimulation-evoked VTA dopamine neuron activity^36^ or dopamine release in the vStr^37^. Disentangling these possibilities will require causal manipulations such as closed-loop SWR disruption and projection-specific inactivation, but either pathway supports the broader conclusion: that hippocampal SWRs/putative replay events are coordinated with the dopaminergic system in a manner suitable for offline learning from experience. These results provide a novel component towards establishing a blow-by-blow account of how individual episodes of internally generated brain activity during SWRs can support learning and decision-making.

## Resource availability

### Lead contact

Requests for further information and resources should be directed to and will be fulfilled by the lead contact, Miriam Janssen.

### Materials availability

This study did not generate new unique reagents.

### Data and code availability

Preprocessed data used to generate the results in this study is available [here](https://drive.google.com/drive/folders/10UE2ELg88IoXdbi_Ajxv_Ak1oMTityr0?usp=sharing). All analyses, described in detail below, were performed using MATLAB 2023a (MathWorks) and can be reproduced using code available through public GitHub repositories (https://github.com/mimijanssen/replayDA, and the van der Meer lab codebase, https://github.com/vandermeerlab/vandermeerlab).

## Acknowledgements

We would like to thank the Neural Systems and Behavior Summer School at Woods Hole for the fruitful discussion and experimental pilot for this research. The authors also thank Dr. Gina Poe and Ward Pettibone who advised us on sleep-stage analysis standard practices.

## Author contributions

H.C., N.T., and M.M. conceptualized the experiments; H.C., M.J., and M.M. were responsible for methodology; M.J. was responsible for data curation, formal analysis and visualization. M.J. and M.M. were responsible for writing.

## Declaration of Interests

The authors declare no competing interests.

## Declaration of generative AI and AI-assisted technologies

During the preparation of this work, the authors used genAI to assist in data organization and visualization. All code was verified and tested by the authors. GenAI was used to detect typographic and formatting inconsistencies in the manuscript text.

## Supplemental information titles and legends

**Extended Figures S1-S3**

## STAR★Methods

### Experimental model and study participant details

#### Study participants

The main data set was collected from eight wild-type mice (C57BL/6J, 3 female, 5 male; 5-10 months old), ranging between 22–31 grams in weight. For the GFP-only study, data was collected from three wild-type mice (C57BL/6J, 3 male, 3-10 months old), approximately 24–33 grams in weight. For the on-track study, data was collected from four additional wild-type mice (C57BL/6J, 3 female, 1 male; 3-6 months old), approximately 19-37 grams in weight.

#### Overview of timeline

mice underwent a single recovery surgery to receive (1) intracranial adeno-associated virus (AAV) injection for the expression of the dopamine sensor GRAB_DA2m_ ^16^, (2) an optic fiber implant to image [DA] in the ventral striatum, and (3) an electrode implant in the ipsilateral hippocampus (dorsal CA1) for the recording of sharp wave-ripples (SWRs). Following recovery from surgery, mice were food-restricted and pretrained in daily sessions to shuttle back and forth on a linear track for liquid rewards before data acquisition began. All procedures were approved by the Institutional Animal Care and Use Committee at Dartmouth College (Dartmouth College IACUC protocol #00002235).

## Method details

### Surgery

Before surgery, mice received a subcutaneous injection of 5 mg/kg ketoprofen and 4 mg/kg dexamethasone. Mice were anesthetized in a closed chamber with 5% isoflurane vaporized in 2L/min medical-grade oxygen. Following induction, mice were transferred to the stereotaxic apparatus, where they continued to receive isoflurane through a nose cone at a concentration of 0.5–2% in 1 L /min oxygen. Mice were shaved to expose the skin over the skull, which was sanitized with betadine and ethanol, and anesthetized locally with lidocaine, before an incision was made. One metal screw was placed in the skull near the future location of the fiber implant to provide stability. A drill burr was used to make a craniotomy above the ventral striatum (from bregma, +1.4 mm anteroposterior and ±0.9–1.0 mediolateral). Mice received two injections of 200 *nL* AAV9-hsyn-GRAB_DA2m_ DA4.4 (WZ Biosciences, YL002009-AV9-PUB)^16^ at 100 *nL* /min unilaterally in the ventral striatum (-3.8 and -4.1 mm dorsoventral from skull surface). GFP-only control mice were unilaterally injected with 200 *nL* AAV8-CamKII-Jaws-KGC-GFP (Addgene, 65015) at the same coordinates. After the viral infusion, a single 400 μm optic fiber mated to a 1.25 mm ferrule was chronically implanted -3.8 mm below the skull surface at the ventral striatal injection site. The screws and ferrule were secured to the skull with dental cement (C&B Metabond, Parkell). A 1.8 mm trephine was then used to make a craniotomy centered above the dorsal CA1 (from bregma, -2.0 mm anteroposterior and ±1.5 mediolateral). Stereotrode microdrives (Axona, described below) were lowered in the craniotomy so that the inner cannulae were implanted 0.2 mm below the pial surface. Two additional ground screws inserted above the ipsilateral cerebellum and contralateral posterior parietal cortex, connected to the microdrive and secured to the skull and microdrive with dental cement. Immediately after surgery, mice were injected subcutaneously with an antibiotic (enrosite; 5 mg/kg) and saline (1 mL). 24h post-surgery, mice were injected subcutaneously with ketoprofen (5 mg/kg), enrosite (5 mg/kg), and saline (1mL); 48h post-surgery, mice were injected with 5 mg/kg of enrosite and 1 mL of saline.

### Behavioral task

Following recovery from surgery (48h minimum), mice received two daily habituation sessions (lap training and exposure to the behavioral apparatus; 5 min each). Mice were then trained to criterion on the behavioral task (described below) until they ran 60 trials in a 30-minute session (training to criterion took 1–6 days). While mice were being trained to criterion, they were placed on food control, with access to approximately 2–5 grams of chow daily and *ad libitum* water until the end of experiments. During food control, weights were maintained approximately above 85% of baseline free-feeding weight (on average 88%). In addition, electrode depths were adjusted to reach the hippocampal cell layer daily following behavioral training.

The behavioral task was run on a 1-meter-long linear track, where running from one end to the other triggered automated evaporated milk delivery using infrared beams. 10% of trials resulted in no reward, 80% in medium volume reward (∼25 μL), and 10% in large volume reward (∼50 μL). Dopamine (DA) fiber photometry signal screening began after at least 9 days of virus expression after surgery. Recordings started once spontaneous [DA] transients and hippocampal SWRs could readily be observed, and once mice reached behavioral criterion on the task (10-31 days post-surgery).

Each daily recording session consisted of a 30 min “pre-task” rest session in the sleep box (a 10 × 10 × 10-inch enclosure with bedding), a ∼30 min task session on a linear track (rewarded, as outlined above), and a 30 min “post-task” rest session in the sleep box. We aimed to record eight sessions for each mouse and four additional amphetamine validation sessions (described below). All data were acquired using a Neuralynx Digital Lynx recording system equipped with Hauppauge HVR USB2 video capture using a ceiling-mounted camera and a custom input stage to route the fiber photometry signal into an unused analog input.

In some sessions, amphetamine was administered to validate the fluorescence signal was modulated by DA. Each of these sessions consisted of a 10-min baseline period preceding the injection of either amphetamine or saline (which alternated daily), followed by a 10 min incubation period. We administered either 0.625–2.5 mg/kg of amphetamine diluted in saline to a volume of 10% the weight of each mouse or just saline at a volume of 10% the weight of each mouse i.p. We adjusted the concentration to have no obvious motor effects while still recording SWRs. Changes in average DA following AMPH administrations were measured as the mean DA from 0 to 11 minutes following injection time, subtracted by the mean DA from 4 minutes prior to injection time.

### Neural data and fiber photometry data acquisition

For hippocampal SWR recordings, we used 16-channel microdrives equipped with a MillMax connector (Axona Ltd). Prior to surgery, a single driveable bundle consisting of 8 stereotrodes (25 *μm* nichrome wire diameter, California Fine Wire Co. RO-800) was cut in a staggered manner, approximately 0.5–1.5 mm in length from the microdrive inner cannula, to provide a depth profile across the hippocampal cell layer for recording SWRs. For fiber photometry, we used Neurophotometrics 0.37 NA black ceramic fibers, fiber diameter 400 *μm*, ferrule diameter 1.25 mm. The fibers were cut to 6.5 mm in length, tested to allow 70% or better light transmission. At the start of each session, implanted microdrives were connected to the data acquisition system using a single Neuralynx HS-18-MM-LED headstage run through a motorized commutator, and signal quality (existence of hippocampal SWRs) checked visually before recording began. For track-mice we used NeuroNexus A1×16-3mm-50-177-CM16LP probes instead of in-house fabricated Axona drives.

GRAB_DA2m_ fiber photometry was performed using a custom photometry system consisting of a photometry cube (FMC3_E(460-490)_F(500-550)_S, Doric) connected to (a) a 470 nm light-emitting diode (M470F3, Thorlabs) via a patch cord (MFP_600/630/-0.48_m_FCM-SMA, Doric), (b) the mouse via a fiber optic patch cable (MFP_400/460/1100-0.48_2m_FCM-MF1.25, Doric or M99L01, Thorlabs) and plastic coupling sleeve (1.25 mm, either from Neurophotometrics or Thorlabs), and (c) a photoreceiver (Model 2151, Newport) via a patch cord (MFP_600/630/-0.48_m_FCM-FCM, Doric). The photodetector output was fed into a DC-coupled analog input of the Neuralynx acquisition system (Cheetah; 5,000 Hz sampling; 1,600 Hz sampling for GFP-only mice). At the start of each session, excitation light intensity was set to 50-100 μW at the tip of the patch cord and the presence of [DA] transients was visually confirmed by streaming a copy of the photodetector output signal using WaveSurfer (Janelia) via a National Instruments ADC board.

## Quantification and Statistical Analysis

### Fiber photometry data acquisition and preprocessing

The fiber photometry signal was first downsampled to 1000 Hz, filtered using a 20 Hz lowpass Butterworth filter, linearly detrended using a 60-second moving window to account for gradual photobleaching (Chronux locdetrend function), and z-scored over the entire session. Signals from GFP-only mice were processed similarly, except that downsampling was omitted. For amphetamine and saline injection sessions the 60 s moving window detrending method was omitted due to the need to measure slow amphetamine-induced changes in the [DA] signal. Instead, we fit a double exponential to the first baseline session (pre-injection) and subtracted that from the signal recorded with amphetamine on board to control for photobleaching while retaining slower changes in [DA].

### Simultaneous electrophysiology data acquisition and preprocessing

Local field potentials (LFPs) were acquired using a Neuralynx Digital Lynx data acquisition system (1500 Hz sampling; 1-500 Hz bandpass filter). Multi-unit activity was recorded by storing a 1-ms snapshot whenever a 32 kHz, 600–6000 Hz filtered signal on any stereotrode channel exceeded an experimenter-set threshold (typically 50 μV). All signals were referenced against ground screws in ipsilateral cerebellum or contralateral posterior parietal cortex.

### Sharp wave-ripple event detection

SWR events were detected based on broad-spectrum frequency similarity to a small set of human-annotated SWRs^35^. For each recording session, the recording channel with the clearest SWRs was identified by the experimenter. On that channel, a noise-corrected SWR Fourier spectrum was obtained by manual identification of example SWR events and subtracting a non-SWR ‘noise’ Fourier spectrum. The dot product between this noise-corrected frequency spectrum and the frequency spectrum of the LFP was obtained using a 60 ms sliding window to obtain a SWR similarity score. Next, we computed a control similarity score by applying the same noise-corrected frequency spectrum dot product to a different recording channel that did not have SWRs; this control similarity score was subtracted from the SWR channel similarity score to obtain an “SWR difference score.” This SWR difference score was then z-scored across the entire session and thresholded at 3-5 standard deviations (SDs, depending on the session) above the mean to obtain candidate SWR events. The resulting candidates were further restricted to SWRs exceeding 20 ms in duration. Detected events that occurred within 20 ms of each other were merged as they were likely from the same SWR. In a separate analysis, we detected SWR events based on thresholded ripple-band power (120-250 Hz; peak z-score > 3 z scores above the mean). The resulting candidates were further restricted to SWRs exceeding 10 ms in duration. Detected events that occurred within 50 ms of each other were merged as they were likely from the same SWR. For on-task analyses, we ran each recording session through DeepLabCut (CITE) to get an estimate of mouse body position over frames and calculated the z-scored speed. If no significant movement (< 0.5 z-score) occurred between 4 seconds prior to 4 seconds after that SWR we kept that SWR for analysis; otherwise it was excluded.

### Sleep-stage labeling

We labeled each two-second segment of our rest sessions as wakefulness or NREM sleep using the <u>AccuSleep GUI</u> (https://github.com/zekebarger/AccuSleep)^36^. Periods of movement were determined as wake periods. Periods of no movement and distinct theta bands were labeled as wake periods. Periods with no movement for longer than 10 seconds with clear delta bands were labeled as NREM periods.

### SWR-triggered (peri-event) DA calculations

For each session, we calculated the average [DA] trace surrounding SWR events (i.e., SWR-triggered DA average, or SWR-DA). To obtain the grand average trace shown in Figure 2a, we took the mean over sessions for each mouse and then took the mean over mice. For the single-session analyses described in the main text, we compared the mean SWR-DA signal for a given session to the distribution of SWR-DA averages obtained from 1,000 resampled sessions in which the DA trace was circularly shifted by a random amount; this resampling procedure results in a permutation z-statistic which we use, first, to determine how many single sessions show significant SWR-DA coupling, and second, as a dependent variable in regressions (described below).

### Permutation tests

#### Amphetamine permutation test

For each mouse, DA traces for all sessions were circularly shifted by a random offset relative to injection time, and a permutation test statistic was calculated as the difference between mock baselined post-amphetamine injection DA and mock baselined post-saline injection DA. This was repeated 1,000 times to generate a null distribution for each mouse. One-sided p values were calculated as the fraction of permutations producing a statistic greater than or equal to the observed difference.

#### Reward prediction error permutation test

For each mouse, DA traces for all sessions were circularly shifted by a random temporal offset relative to reward zone entry times, and a permutation test statistic was calculated as the difference in mean DA within 4 seconds between mock high-volume trials and omission trials. This was repeated 1,000 times to generate a null distribution for of the test statistic for each session. The observed statistic (the difference in mean DA activity during the same 4-second window between the true high-volume and omission trials) was then compared with the null distribution. A one-sided permutation p value was calculated as the fraction of permutations producing a statistic greater than or equal to the observed statistic.

### Session-based regression analyses

For this analysis we treat each rest period (pre-task and post-task “session”) as a data point, meaning there will be two sessions for each recording day (n = 92). To assess whether SWR-triggered DA increases relative to the pre-SWR baseline, we ran session-based mixed-effects regressions of the following form:

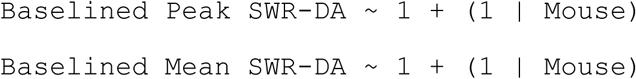

where the dependent variable `Baselined Peak` or `Mean SWR-DA` denotes either the peak or mean (in the 1 second following the SWR) of the baselined, session-averaged SWR-DA signal. Specifically, the session-averaged SWR-triggered PETH was baselined by the mean -4 to -1 seconds before SWR time. The actual dependent variable is the peak (or mean) of this session-averaged DA signal one second following SWRs subtracted by the peak (or mean) of the averaged one second DA signal before SWRs. A positive intercept in this model indicates baselined SWR-DA is greater than zero, i.e. an increase relative to baseline.

To assess whether there was a relationship between SWR-DA strength and (1) the strength of putative DA reward prediction errors (measured during the task; Figure 1d) or (2) motivation, as measured by average running speed on the track, we ran session-based mixed-effects regressions of the following forms:

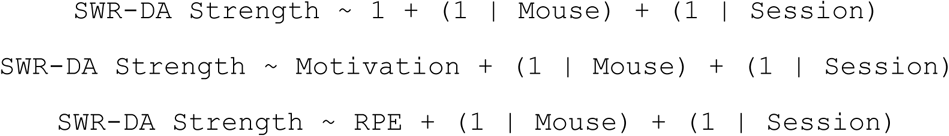

Where `SWR-DA strength` denotes the permutation z-score of the SWR-DA coupling as described in the previous section “SWR-triggered (peri-event) DA calculations” (i.e. raw SWR-DA compared to a circular shuffle for each session). `Motivation` was defined as the average track speed (pixels/s). `RPE` denotes the t-statistic between the mean DA signal for high reward trials and the trough for omission trials within a four second period after reward zone entry.

### Event-based regression analyses

For this analysis we treat each individual SWR as a data point. To assess whether SWR-triggered DA exceeds the pre-SWR baseline, we ran event-based mixed-effects regressions of the following form:

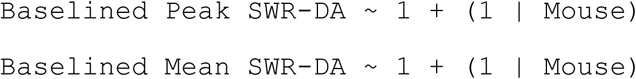

where the dependent variable was obtained by first baselining each individual SWR’s peri-event DA signal (mean DA from -4 to -1 seconds relative to SWR time). Then, for each event, we took the peak (or mean) within one second following a SWR minus the peak (or mean) within one second before SWR. A positive intercept in this “base” model indicates baselined SWR-DA is greater than zero, i.e. an increase relative to baseline.

To examine whether specific SWR properties influence SWR-triggered DA, we ran additional event-based LMMs. We compared the base model above with a model with one additional independent variable:

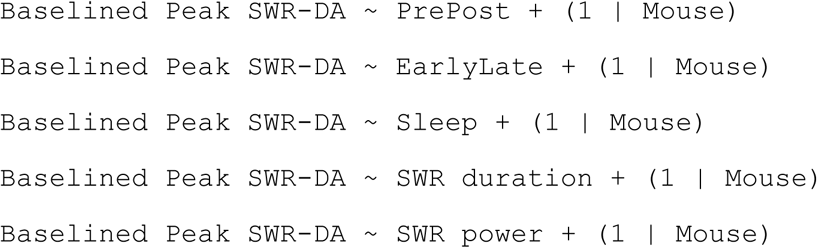

where the independent variable PrePost indicates a categorical variable 1 for pre- and 2 for post-track rest sessions. For track-mice, PrePost indicates a categorical variable 1 for pre-track rest, 2 for track, and 3 for post-track rest sessions. The independent variable EarlyLate indicates a categorical variable 1 for sessions 1-4 and 2 for sessions 5-8. The independent variable Sleep indicates a categorical variable 1 for wake periods and 2 for NREM periods. The independent variable SWR duration indicates the duration of the SWR in milliseconds. The independent variable SWR power indicates the z-scored SWR power within the frequency range 120-250 Hz.

We then compared these models to the base model using simulated likelihood ratio tests. To test for covariance, we also ran simulated likelihood ratio tests between a full model (all dependent measures) against the same model with one dependent measure left out. The full model was of the following forms:

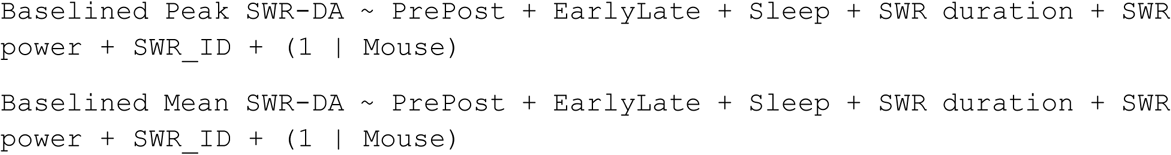

To examine if SWR order had an effect, SWR_ID indicated the SWR number within a session normalized by how many SWRs were detected. We also ran similar models using Baselined Mean SWR-DA as the dependent measure.

### Quantification and statistical analyses

For Figures 1 and Figure 2 and associated analyses, we treated individual sessions as the unit of analysis, and plots of aggregated data show mean +/- standard error of the mean (SEM) over mice. Only sessions in which fiber recording locations were histologically verified to be within the ventral striatum, had power spectral density plots free from noise, raw fiber photometry amplitudes > 0.01 V, at least 100 detected SWRs, sufficiently motivated mice, and no recording failures were included in the analysis (For main experimental mice, N = 92 rest sessions, i.e. 46 recording days, across 8 mice; for GFP mice, N = 48 rest sessions, i.e. 24 recording days, across 3 mice), with a minimum of 4 recordings per mouse. We had to exclude mice with fiber locations outside of the ventral striatum (1 mouse) or if GRAB_DA2m_ was not well expressed (2 mice; none of these were included in the 8 mice analyzed here). Additionally, we excluded recording sessions due to recording failures, such as an optical fiber becoming unplugged (1 recording session). Additionally, sessions which had large noise in the power spectral density plots (6 recording sessions), in which raw fiber photometry amplitudes were below 0.01 V (1 recording session), or in which less than 100 sharp-wave ripples were detected (3 recording sessions) were excluded. Recording days in which the animal did not complete 60 trials in under 30 minutes were also excluded (1 recording session).

For Figure 3 and associated analyses, we treated individual SWRs as the unit of analysis, and plots of aggregated data show the mean without standard deviation. To allow for a better visual comparison of the averaged peri-event time histogram we wrote the mean and standard deviation in the figure caption rather than plotting the standard deviation. This plot also included track mice, which included 96 sessions (pre-track rest, track, and post-track rest), which amounts to 32 recording days across 4 mice.

## Supplemental Figures

**Extended Figure 1.**
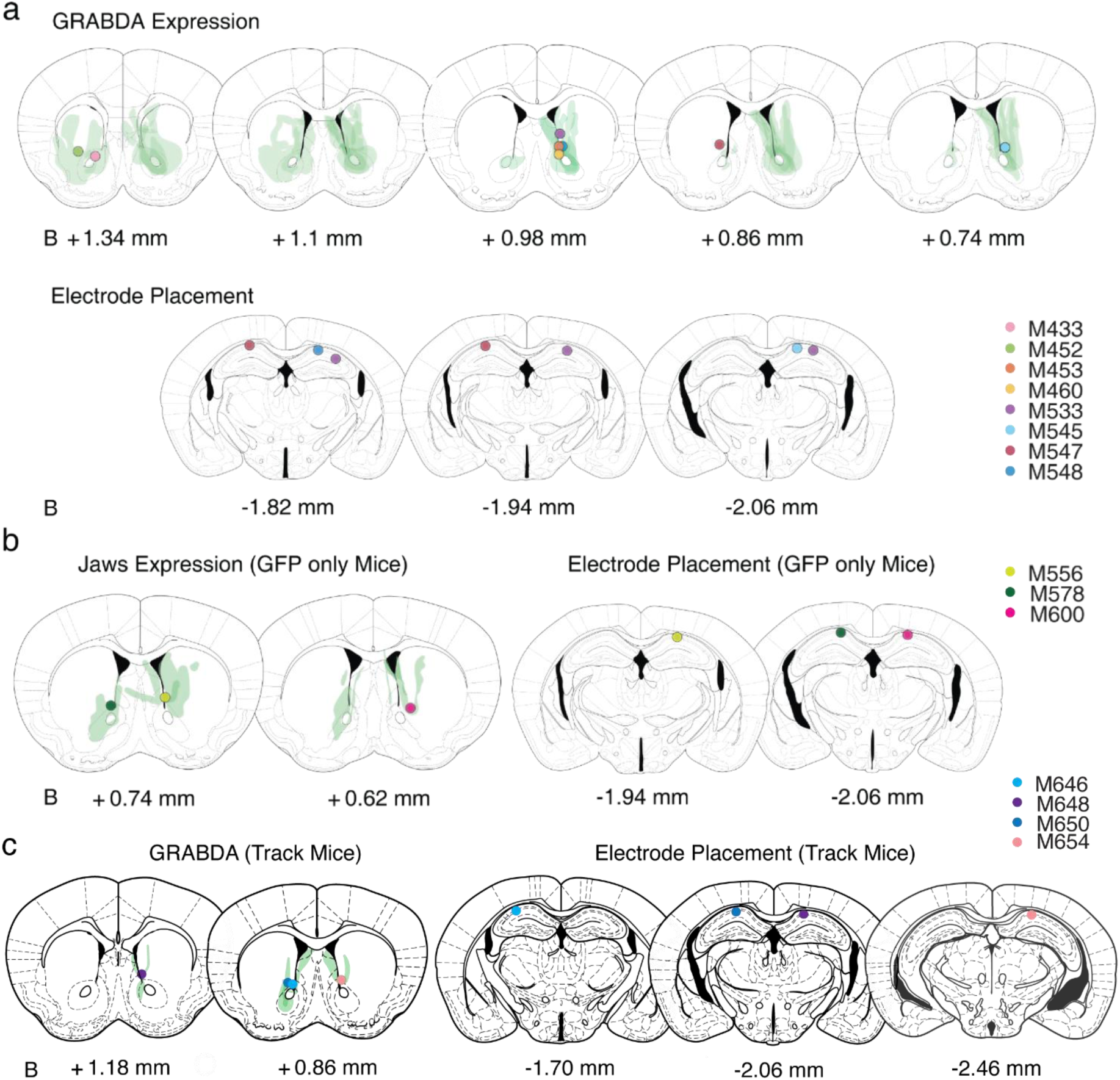
Histological verification of GRAB_DA2m_ expression, GFP-only expression, and microdrive placement. **a**, *Top*: GRAB_DA2m_ expression maps. Each coronal section represents a section relative to bregma (B, in mm). Expression for each animal is plotted at 20% transparency with per-animal expression overlaid. The estimated widest part of the fiber cannula is depicted by a filled circle marker with each color representing a different mouse. *Bottom*: estimated microelectrode placement, by a filled circle marker (same as above) determined by gliosis induced by electrolytic lesions. Three animals did not have clear gliosis, but did have neurophysiological signatures indicating dCA1 placement (Extended Figures 2, 3) as well as clear microdrive cannula placement in the cortex above dCA1. **b**, as in a, for GFP-only mice. Expression for each animal is plotted at 33% transparency with per-animal expression overlaid. **c**, as in a, for track mice. Expression for each animal is plotted at 40% transparency with per-animal expression overlaid.

**Extended Figure 2.**
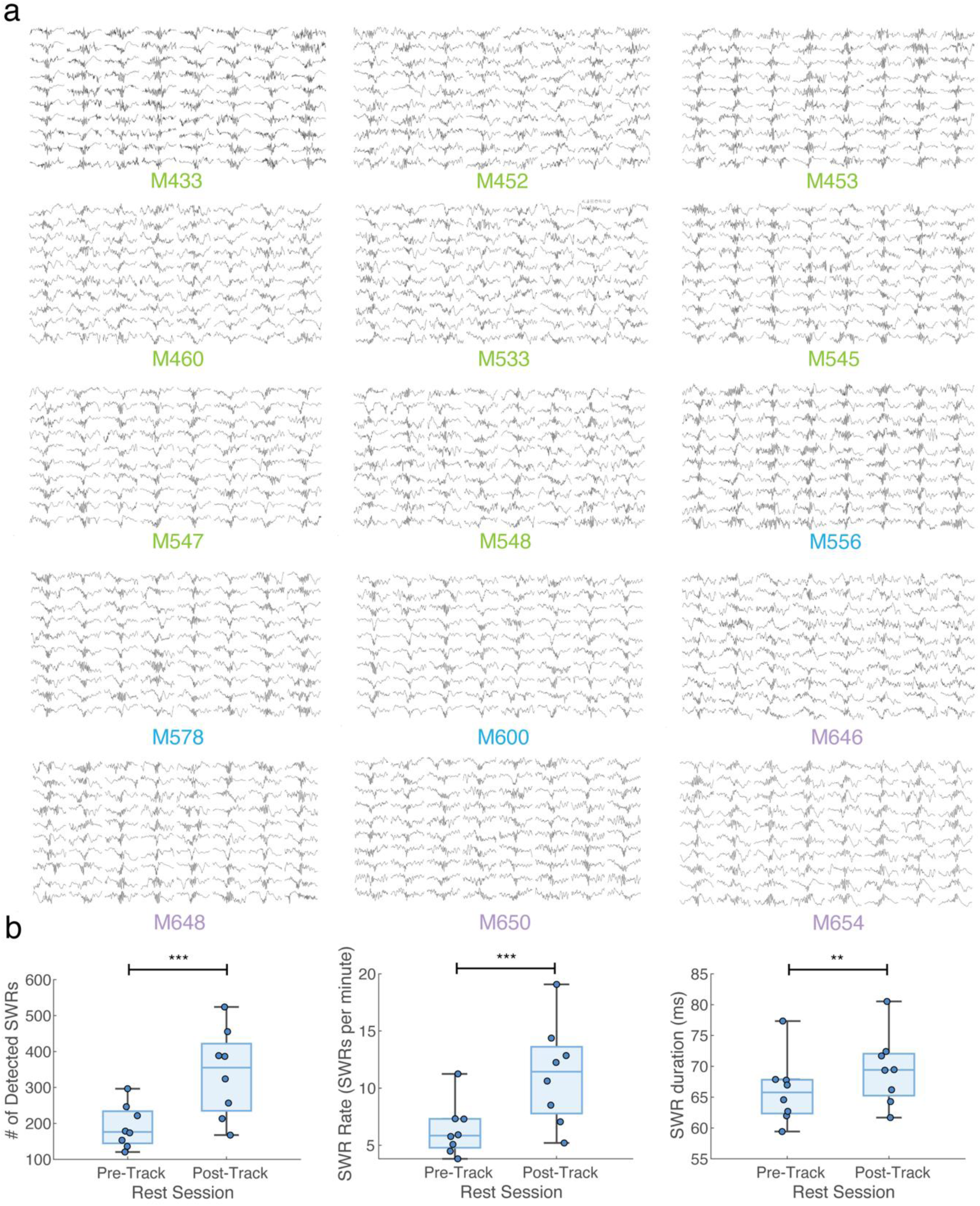
Example SWR traces from each mouse, and sharp wave-ripple descriptive statistics. **a,** 80 example detected SWRs from a session for each GRAB-DA mouse (green), GFP mouse (blue), and track mouse (purple). We consistently observed clear SWR events in each included mouse, with no visible differences between GRAB-DA and GFP-only mice. **b,** *Left*: Average SWR count. The average number of detected SWRs, separated by pre- and post-task rest sessions. Each marker represents an average over sessions for a mouse; whiskers indicate the maximum and minimum. The average number of SWRs was 190.86 ± 59.79 SWRs for pre-task rest sessions and 339.56 ± 122.30 SWRs for post-task rest sessions (paired t-test, t(7) = -5.72, p = 7.17e-04). *Middle*: Average SWR rate: 6.60 ± 2.12 SWRs per minute for pre-task rest sessions and 11.81 ± 3.78 SWRs per minute for post-task rest sessions (paired t-test, t(7) = -7.06, p = 2.01e-04). *Right*: SWR duration: 66.23 ± 5.26 ms for pre-task rest sessions and 69.72 ± 5.41 ms for post-task rest sessions (paired t-test, t(7) = -4.88, p = 0.0018). *** indicates p < 0.001, ** indicates p < 0.01, * indicates p < 0.05.

**Extended Figure 3.**
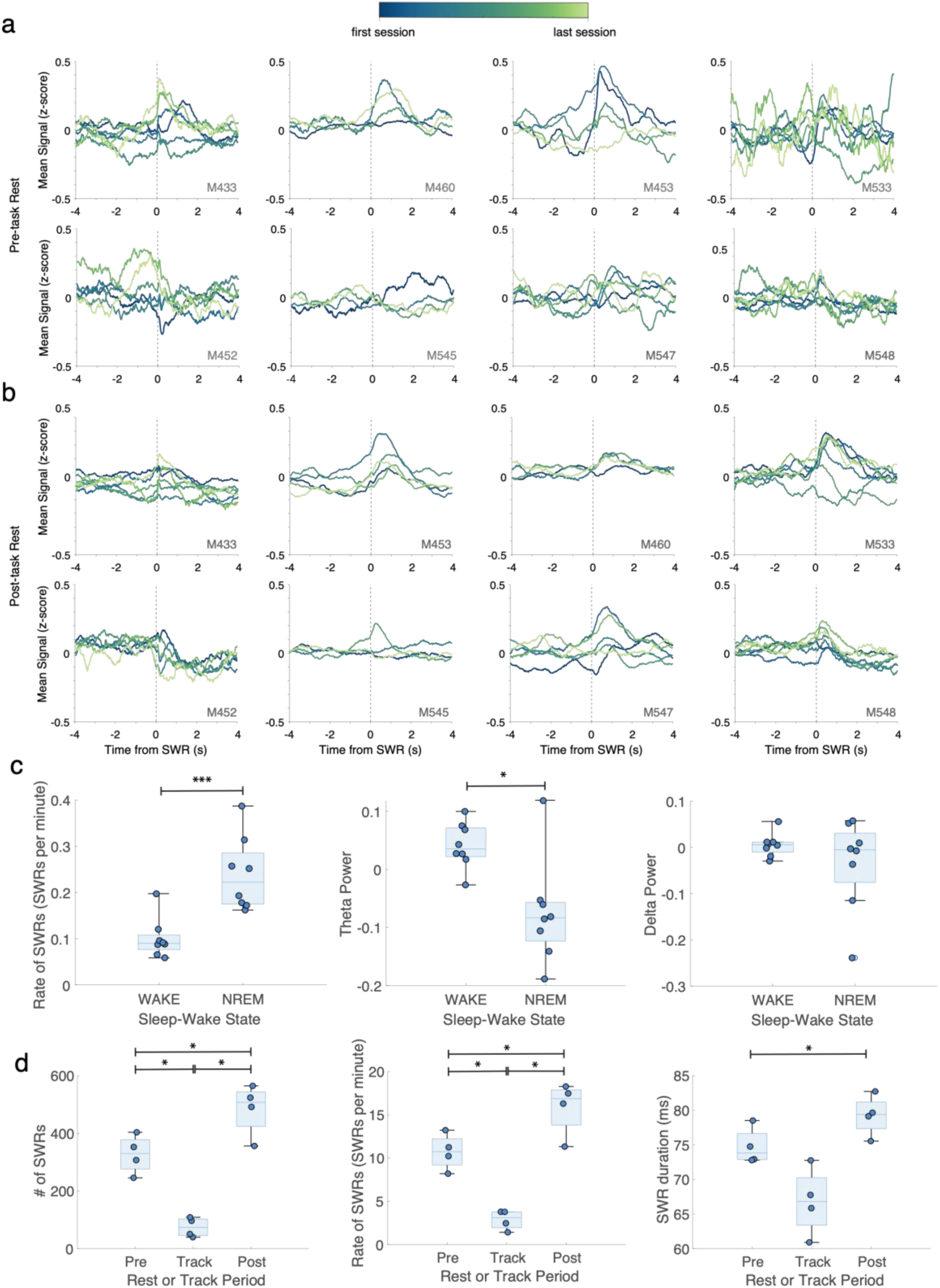
Peri-event time histograms for SWR-triggered dopamine response for each individual session, and further SWR properties. **a,** Average SWR-triggered DA for each pre-task rest session. Sessions are graded in color with the first session in dark blue and the last session for that mouse in light green. **b,** as in *a*, for post-task rest sessions. **c**, *Left*: Average SWR rate, separated by wake and NREM periods. Each marker represents an average over sessions for a mouse; whiskers indicate maximum and minimum. The average rate of SWRs was 0.10 ± 0.043 SWRs per second for wake periods and 0.24 ± 0.080 SWRs per second for NREM periods (paired t-test, t(7) = - 7.98, p = 9.27e-05). *Middle*: Average theta power separated by wake and NREM periods: 0.041 ± 0.040 z-score power for wake periods and -0.075 ± 0.090 z-score power for NREM periods (paired t-test, t(7) = 2.69, p = 0.031). *Right*: Average delta power separated by wake and NREM periods: 0.0050 ± 0.025 z-score power for wake periods and -0.035 ± 0.099 z-score power for NREM periods (paired t-test, t(7) = 0.94, p = 0.38). *** indicates p < 0.001, ** indicates p < 0.01, * indicates p < 0.05. **d,** *Left*: Average SWR count for the track cohort. The average number of detected SWRs, separated by pre-task rest, track, and post-task rest sessions. Each marker represents an average over sessions for a mouse. The whiskers are the maximum and minimum. The average number of SWRs was 327.4 ± 67.1 SWRs for pre-task rest sessions, 74.2 ± 33.9 SWRs for track sessions, and 484.1 ± 90.3 SWRs for post-task rest sessions. We found a significant difference in SWR count between pre-task rest and track sessions (post-hoc paired t-test, t(3) = 5.76, p = 0.0104), between pre- and post-task rest sessions (post-hoc paired t-test, t(3) = - 6.35, p = 0.0079), and between track and post-task sessions (post-hoc paired t-test, t(3) = -8.042, p = 0.0040). *Middle*: Average SWR rate for the “track” cohort: 10.73 ± 2.09 SWRs per minute for pre-task rest sessions, 2.87 ± 1.15 SWRs per minute for track sessions, and 15.85 ± 3.12 SWRs per minute for post-task rest sessions; significant difference between pre-task rest and track sessions (paired t-test, t(3) = 5.17, p = 0.0104), between pre- and post-task rest sessions (paired t-test, t(3) = -5.82, p = 0.0101), and between track and post-task sessions (paired t-test, t(3) = -6.92, p = 0.0062). *Right*: Average SWR duration for the track cohort, separated by pre-task rest, track, and post-task rest sessions. The average SWR duration was 74.75 ± 2.67 ms for pre-task rest sessions, 66.83 ± 4.90 ms for track sessions, and 79.27 ± 2.94 ms for post-task rest sessions. SWR duration was not different between pre-task rest and track sessions (paired t-test, t(3) = 2.76, p = 0.0704), or between track and post-task rest (paired t-test, t(3) = -3.48, p = 0.0401) but was different between pre- and post-task rest sessions (post-hoc paired t-test, t(3) = -5.17, p = 0.0140). Bonferroni-corrected alpha levels: *** indicates p < 0.00033, ** indicates p < 0.0033, * indicates p < 0.0167.

## Notes

Funding: This work was supported by the National Institute of Mental Health grant NIH R01 MH123466 (to MvdM) and the National Science Foundation NSF GRFP 2236868 (to MJ).

### Competing Interest Statement

The authors have declared no competing interest.

### Summary of Updates

- added new cohort of subjects, replicating original result and extending results to contain comparison of on-task vs off-task rest periods - included many new statistical analyses

